# The C-terminal ZZ domain of the Drosophila ORB2 RNA-binding protein is required for spermatid individualization

**DOI:** 10.1101/2025.08.22.671863

**Authors:** Timothy C.H. Low, Brook L. Falk, Julie A. Brill, Howard D. Lipshitz

## Abstract

ORB2 is the Drosophila ortholog of the human CPEB2-4 family of RNA-binding proteins, which include a conserved C-terminal Zinc-binding (‘ZZ’) domain. We have recently shown that this domain interacts with several translation co-repressors in the early embryo, and that deletion of this domain from the endogenous *orb2* gene results in derepression of its target mRNAs (LOW *et al*. 2025). Here we assess the effect of deletion of the ZZ domain on spermatogenesis. We find that ORB2ΔZZ protein is no longer localized to the distal tip of spermatids; that in *orb2^ΔZZ^* testes, additional proteins known to play a role in spermatid individualization – ORB, IMP, SOTI – are mislocalized; that the SOTI-dependent Cleaved Caspase 3 gradient no longer forms; that individualization complexes are defective; and that *orb2^ΔZZ^* flies are sterile and lack mature sperm.

## Introduction

Proteins belonging to the cytoplasmic polyadenylation element binding (CPEB) family are highly conserved RNA-binding proteins that canonically interact with U-rich cytoplasmic polyadenylation element (CPE) motifs found in the 3ʹUTR of target transcripts (HAKE AND RICHTER 1994; IVSHINA *et al*. 2014). Most animals, including humans, mice, and *Caenorhabditis elegans* have four CPEB genes (where CPEB2-4 are more closely related to each other than to CPEB1), whereas *Drosophila* has two: *orb* and *orb2*. While the N-terminal portion of CPEB proteins is divergent, there are high levels of homology at the C-terminus of CPEB proteins, mostly concentrated at the tandem RNA-recognition motifs (RRMs) that serve as the RNA-binding domain (RBD) and its adjacent ZZ-class zinc binding domain.

CPEB2-like proteins are expressed in both the soma and the germline, with functions in the nervous system, asymmetric stem cell division, and gametogenesis (IVSHINA *et al*. 2014). In *Drosophila*, *orb2* is enriched in the testis, where it is essential for many stages of spermatogenesis (XU *et al*. 2012; VEDELEK *et al*. 2018; LI *et al*. 2022). The *orb2* locus produces two protein isoforms, and both are expressed in the testes; the 60-kDa ORB2A and the 75-kDa ORB2B isoforms share an identical C-terminal sequence containing 542 amino acids, and differ only at their N-termini, which contain 9 and 162 unique amino acid sequences, respectively.

In the testis (diagrammed in Figure 1), ORB2 is detectable after the completion of mitosis in the 16-cell cysts, where expression of both *orb2* mRNA and protein rapidly increase and peak during meiosis, specifically at the 32 and 64 cell spermatid cysts (XU *et al*. 2012). After this stage, ORB2 persists and is distributed along the entire axoneme bundle, with the highest concentration found at the distal tip of the growing flagellar axonemes. This expression pattern corresponds with the phenotype of *orb2* null mutants, in which spermatocytes fail to complete meiosis and have defects in spermatid differentiation; these disruptions cause male sterility (XU *et al*. 2012; XU *et al*. 2014). In addition, deletion of the *orb2* mRNA’s 3’UTR has no effect on meiosis, but results in defects in spermatid individualization that may be attributable to mislocalization of *orb2* mRNA and ORB2 protein in the 64-cell cyst (GILMUTDINOV *et al*. 2021).

**Figure 1.**
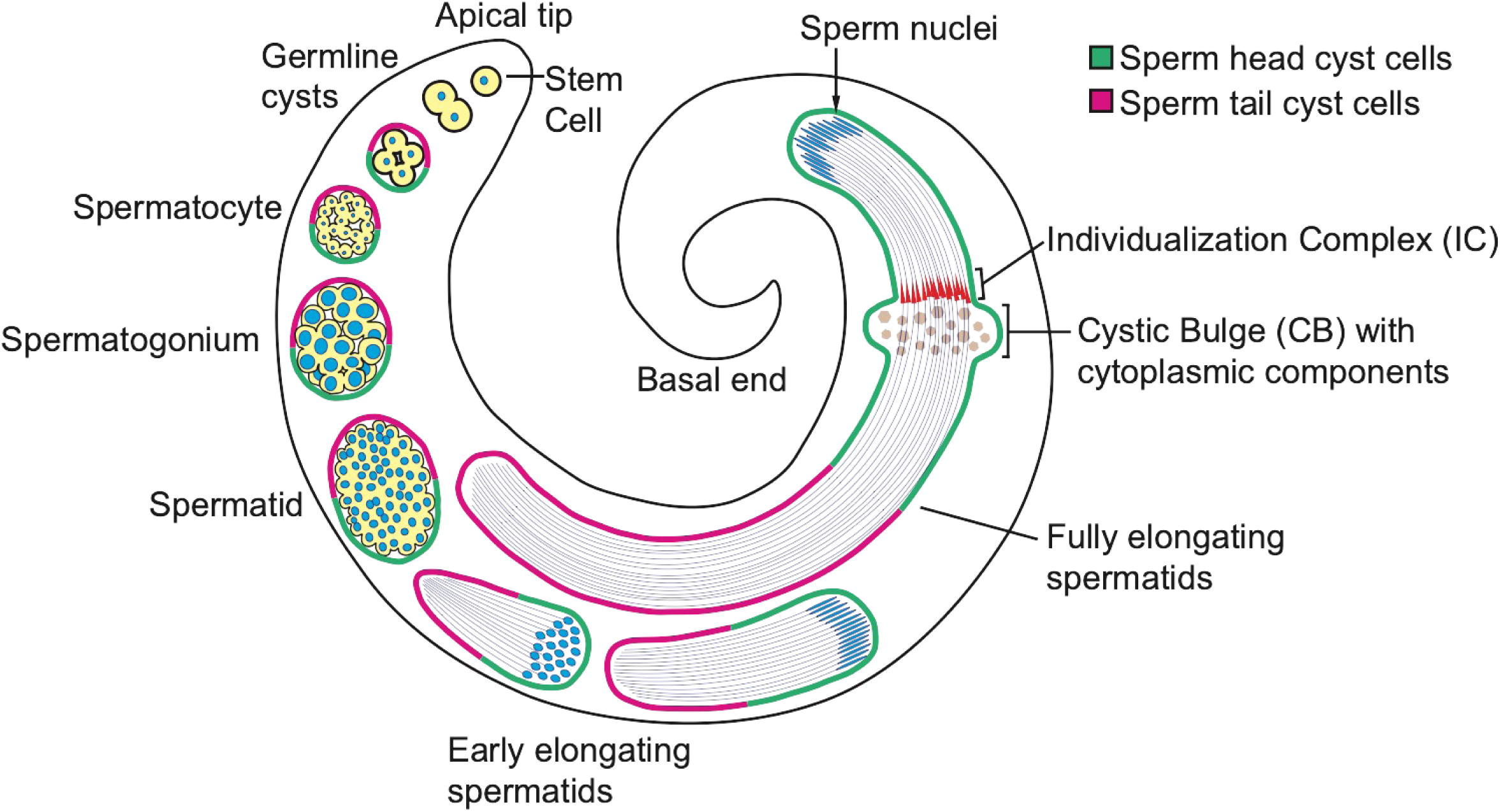
Diagram depicting the stages of spermatogenesis as they are spatiotemporally organized in the Drosophila testis. Germline cysts are oriented with posterior/head cyst cells (green) pointing towards the basal tip, whereas the anterior/tail cyst cells (magenta) are pointing towards the apical tip. Cysts travel from the apical tip towards the basal tip as they mature.

We recently reported that the ORB2 protein interacts with hundreds of target transcripts and physically interacts with multiple translational repressors in early embryos (LOW *et al*. 2025). An endogenous deletion of the ZZ domain (*orb2^ΔZZ^*) produced by CRISPR, which retains the endogenous RBD and 3’UTR, binds to target RNAs but loses interaction with many of the co-repressors, resulting in derepression of target mRNA translation in embryos during the maternal-to-zygotic transition (LOW *et al*. 2025). *orb2^ΔZZ^* females are fertile and their embryos have no overt developmental phenotypes.

Here, we show that male *orb2^ΔZZ^* flies are sterile and we assess the role of the ZZ domain during spermatogenesis in the adult male testis. We show that the ZZ domain has a minor role during meiosis but is essential for spermatid individualization, failure of which leads to male sterility in *orb2^ΔZZ^* mutants. Loss of the ZZ domain results in increased levels of ORB2 but does not affect its spatial expression in the testis. In *orb2^ΔZZ^* mutant testes, individualization complexes (ICs) are mispositioned and actin cones are scattered. Moreover, ORB, IMP, SOTI and Cleaved caspase 3 (CC3) are mislocalized. Our data are consistent with a model in which the ZZ domain of ORB2 is required for organization of the distal region of the spermatid cyst and establishment of the SOTI gradient, which in turn regulates CC3 production and successful spermatid individualization.

## Materials and Methods

### Fly stocks

The following *Drosophila melanogaster* stocks were used: *w^1118^*is the host, pre-injected *w^1118^* line that was used to generate the CRISPR *orb2^ΔZZ^* mutant line (LOW *et al*. 2025). Mutants analyzed were *orb2^ΔZZ^* (LOW *et al*. 2025), *orb2^36^* and *orb2^7^*(XU *et al*. 2012). Adult *orb2^ΔZZ^*/*orb2^36^* and *orb2^ΔZZ^*/*orb2^7^*flies were obtained by crossing homozygous *orb2^ΔZZ^* virgin females with *orb2^36^*/balancer or *orb2^7^*/balancer males. The methods used to generate *orb2^ΔZZ^* are described in (LOW *et al*. 2025); *orb^36^* (BDSC#58479) and *orb2^7^* (BDSC#58480) were obtained from the Bloomington Drosophila Stock Center, deposited by Paul Schedl. Details of the stocks can be found on FlyBase (OZTURK-COLAK *et al*. 2024). Flies were cultivated at 25°C under standard laboratory conditions unless otherwise indicated.

### RT-qPCR

RNA was isolated from 1-3 day post-eclosion adult testes using TRI Reagent (Sigma) according to manufacturer’s instructions. Isolated RNA was quantitatively reverse transcribed into single-stranded cDNA with the Superscript IV reverse transcriptase kit (Invitrogen) according to manufacturer’s instructions. 500ng of total RNA per sample was used to synthesize cDNA, which was primed using random hexamer primers. The resulting single-stranded cDNA was diluted 1:25 using RNase-free water and used to perform quantitative real-time PCR with primers specific to various transcripts assayed as derived from FlyBase (Figure S1) (OZTURK-COLAK *et al*. 2024). Primers specific to the transcripts assayed were designed using NCBU Primer-BLAST. Qualitative real-time qPCR was performed with SensiFAST SYBR PCR mix (Bioline) following the manufacturer’s protocol and using 5 µL of diluted cDNA per reaction.

CFX384 Real-Time System (Bio-Rad) was used to carry out the PCR reaction.

The results of the qPCR were analyzed using CFX Manager software (Bio-Rad). Each biological replicate represents a measurement obtained for a separate set of testis dissections; values from three technical replicates are averaged and relative gene expression was normalized to the ribosomal protein-coding transcript *RpL32* and *GAPDH2* as normalization controls. Normalized expression is presented as the average of three biological replicates with error bars representing standard deviation calculated by one-way ANOVA.

### Antibodies

The following primary antibodies were used: mouse anti-ORB2 4G8 (1:50 for Western blot, 1:200 for immunostaining; obtained from the Developmental Hybridoma Bank), mouse anti-ORB 6H4 (1:200 for immunostaining; obtained from the Developmental Hybridoma Bank), rabbit anti-IMP (1:100 for immunostaining; gifted by Paul Macdonald), Guinea pig anti-SOTI (1:50 for immunostaining; gifted by Eli Arama), rabbit monoclonal anti-Cleaved Caspase 3 (Asp175, Cell Signaling Technology). Mouse anti-Actin (Sigma) was used at 1:10,000 for Western blots as a loading control. The following secondary antibodies were used in the immunostaining: goat anti-mouse Alexa555, goat anti-mouse Alexa488, goat anti-rabbit Alexa555, goat anti-rabbit Alexa488, and donkey anti-Guinea pig Cy3.

### Western Blots

Proteins were resolved by SDS-PAGE and transferred to PVDF membrane, blocked at room temperature for 30 minutes with 1% non-fat milk in PBST (1xPBS supplemented with 0.1% Tween20). Blots were hybridized with primary antibodies in 1% milk in PBST at 4°C overnight while nutating. HRP-conjugated secondary antibodies were hybridized to the blot at 1:5000 dilution in 1% milk in PBST at 20°C while nutating for 1 hour. Blots were developed using ECl detection substrate (Millipore Immobilon Luminata Crescendo Western HRP substrate) and imaged with ChemiDoc and ImageLab (BioRad).

### Male Fertility Assay

Male flies were collected and individual males were crossed with two *w^1118^* virgin females in a vial with yeast pellets for 7 days; the adults were removed, and the vials were scored for the presence of larvae, pupae, and adults for the following 7 days. Males whose matings produced larvae were scored as fertile.

### Testis Immunostaining

For whole mount α-ORB2, α-IMP, α-ORB, and α-SOTI staining (Figures 2, 6-8), testes from 1-to 3-day old flies were dissected into 4°C PBS and fixed with 1% paraformaldehyde for 20 minutes. After fixation, testes were washed three times with PBST (1x PBS + 0.1% Tween20) for 5 minutes (here and below). Testes were then washed twice with PBST and incubated with a solution of 0.1% Tween-20 and 0.3% Triton X-100 in 1×PBS) with 5% normal goat serum (Life Technologies) at room temperature for 1 hour, which was followed by an overnight incubation with primary antibody and/or rhodamine phalloidin. This was followed by three washes with PBST and incubation with secondary antibody for 2 hours. After three final washes with PBST, testes were mounted on slides with DAPI in DABCO mounting media. Images were acquired on an inverted Leica DMi8 epifluorescence microscope with a 40x phase-contrast objective and Leica K5 camera using the Thunder Imaging System.

**Figure 2.**
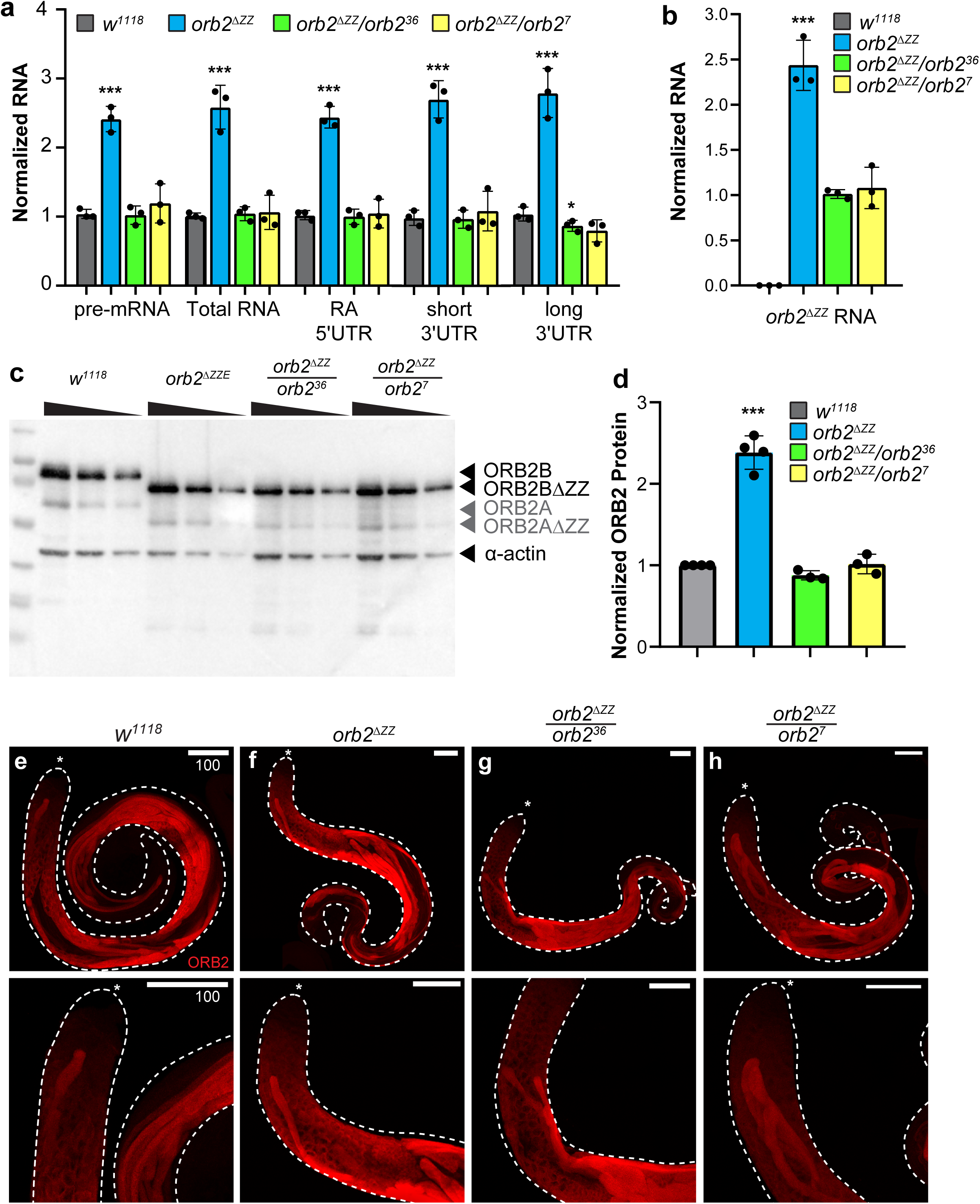
Expression of *orb2* mRNA and ORB2 protein in testes. (a) and b) Bar graphs showing the expression of various *orb2* mRNA isoforms (primer targets shown on the x-axis and diagrammed in Figure S1) measured by RT-qPCR from testis extract prepared from four different genotypes (indicated by bar color). RNA levels shown are double normalized, first to *RpL32* and *GAPDH2* mRNAs, and then to *w^1118^*. c) Western blot showing the expression of ORB2 and ORB2ΔZZ protein in testis extracts. d) Bar graph showing the quantification of ORB2 and ORBΔZZ protein levels normalized to β-tubulin loading control; n=4. e-h) Confocal microscope images of testes stained with anti-ORB2 antibody in the genotypes indicated. Asterisk (*) marks the apical tip of the testis; scale bars represent 100μm.

For Hoechst staining (Figure 4), testes were dissected in cold PBS with Hoechst-33342 (1:1000) and transferred onto poly-L-lysine coated slides with a drop of PBS and Hoechst (as above). The apical tip of each testis was cut off, a coverslip was placed on top, and excess PBS was removed with a Kimwipe to draw cells out of the testis sheath. Images were acquired on an inverted Leica DMi8 epifluorescence microscope with a 40x phase-contrast objective and Leica K5 camera using the Thunder Imaging System. Images were uniformly processed for brightness and contrast using LAS-X software.

For α-CC3 (Figure 9), 1-to 3-day old testes were dissected in testis isolation buffer (183 mM KCl, 47 mM NaCl, 10 mM Tris pH 6.8) and transferred onto a poly-L-lysine coated slide.

Coverslips were placed over the testes and flattened by drawing off liquid with a Kimwipe. Slides were frozen in liquid nitrogen for 10 seconds before the cover slip was removed with a razor blade. Slides were incubated in-20°C 95% ethanol bath for 10 minutes. Samples were fixed in 4% paraformaldehyde for 7 minutes, washed in PBS + 0.1% Triton X-100, permeabilized in PBS + 0.3% Triton X-100 + 0.3% Na Deoxycholate for 15 minutes twice, washed for 10 minutes, and blocked in PBS + 0.1% Triton X-100 + 5% Bovine Serum Albumin for 30 minutes. Samples were incubated overnight in blocking solution with α-CC3 (1:400), washed thrice for 5 minutes and once for 15 minutes, and incubated in secondary antibody for one hour. Samples were washed for 15 minutes, incubated for 15 minutes in PBS + 0.1% Triton X-100 with DAPI (1:1000) and 30 minutes in PBS + 0.1% Triton X-100 with rhodamine-phalloidin (1:200). Following this, samples were washed twice for 15 minutes and mounted with fluorescence mounting medium. Images were acquired on an inverted Nikon A1R laser scanning confocal with 10x and 60x objectives using NIS Elements software. Images were uniformly processed for brightness and contrast using Volocity software. Experimental replicates were carried out with the following modifications: testes were dissected in cold PBS and transferred into membrane-lined trays in four-well plates where they were fixed for 30 minutes, permeabilized for 30 minutes in PBS + 0.3% Triton, blocked in PBS + 0.3% Triton + 0.5% BSA, and mounted onto poly-L-lysine coated slides in Vectashield Antifade Mounting Medium with DAPI. Similar phenotypes were observed with both protocols.

## Results

### Deletion of the ZZ domain results in increased expression of orb2 mRNA and ORB2 protein in testes

To determine whether deletion of the ZZ domain affects expression of the *orb2* gene in testes, we examined levels of transcripts in testis extracts by RT-qPCR and protein by Western blot (Figure 2). RT-qPCR of *orb2* mRNA in testis extracts showed that *orb2^ΔZZ^*transcripts were present at roughly double the level in *orb2^ΔZZ^*homozygous testes compared to WT *orb2* transcripts in *w^1118^*testes (Figures 2a and S1). *orb2^ΔZZ^* mRNA levels were comparable to *orb2* levels in wild type when *orb2^ΔZZ^* was present in a single copy in combination with either of two previously reported *orb2* deletion mutants – *orb2^36^* and *orb2^7^* – which each harbor a different targeted deletion of the *orb2* locus (Figure 2a) (XU *et al*. 2012). To ensure that the *orb2^ΔZZ^* transcripts contained the precise ZZ domain deletion, we also used a pair of primers that span the deletion site to amplify and quantify the levels of *orb2^ΔZZ^* transcript in testis extract from each genotype (Figure S1). As expected, no PCR amplification of *orb2^ΔZZ^*transcript was detected in *w^1118^* testes (Figure 2b), while expression levels in the *orb2^ΔZZ^* homozygote and *orb2^ΔZZ^*/deletion hemizygotes were comparable to that shown in Figure 2a.

We next examined protein expression and found that, consistent with our analyses of mRNA, there was two-fold more ORB2ΔZZ protein in *orb2^ΔZZ^*testes than ORB2 protein in wild-type testes (Figure 2c,d). Also consistent with the mRNA results, ORB2ΔZZ protein levels in testes from *orb2^ΔZZ^*/deletion heterozygous flies were comparable to ORB2 protein levels in wild-type testes (Figure 2c,d).

Over-expression of *orb2^ΔZZ^* mRNA and ORB2ΔZZ protein is likely due to loss of negative autoregulation, since ORB2 is known to regulate expression of its own mRNA (MASTUSHITA-SAKAI *et al*. 2010; STEPIEN *et al*. 2016; LOW *et al*. 2025).

### Deletion of the ZZ domain does not affect spatial expression of ORB2 in testes

To determine whether the expression pattern and localization of ORB2 protein in the testis is affected by the ZZ domain deletion, we stained 3-day old testes with mouse anti-ORB2 antibodies and examined the expression of ORB2 protein by laser scanning confocal microscopy (Figure 2e-h). Consistent with previous reports (XU *et al*. 2012; GILMUTDINOV *et al*. 2021), in *w^1118^* testes there was little to no ORB2 protein in the stem cells and low levels began to be detected in early primary spermatocytes (Figure 2e). After completion of mitosis, as the spermatogonia transitioned through S-phase into primary spermatocytes, there was a significant increase in ORB2 protein corresponding with the extended G2 phase associated with high levels of transcription and stockpiling of gene products required for subsequent processes. ORB2 protein persisted in the post-meiotic 64-cell cysts as they underwent the process of spermatid elongation, where it was distributed along the entire length of the sperm tails, with a slight enrichment of ORB2 protein in the distal end of elongated spermatid cysts (i.e., the opposite end to the nuclei) (Figure 2e).

In the *orb2^ΔZZ^* homozygous mutant testes, while the level of ORB2ΔZZ was increased as described (Figure 2c,d), the spatial expression pattern was indistinguishable from *w^1118^* testes (Figure 2f). We also examined the expression of ORB2ΔZZ protein in testes dissected from *orb2^ΔZZ^*/deletion heterozygotes and, again, found that the expression pattern of ORB2ΔZZ protein was comparable to that in *w^1118^* testes (Figures 1g, h).

Taken together, these results indicate that the timing and expression pattern of the ORB2ΔZZ protein is similar to that of wild-type ORB2 protein during spermatogenesis.

### orb2^ΔZZ^ males are sterile and do not produce mature sperm

Deletion of the *orb2* gene (XU *et al*. 2012) or its 3’UTR (GILMUTDINOV *et al*. 2021) causes male sterility. To determine whether deletion of the ZZ domain had any effect on male fertility, we performed a fertility assay on adult male *orb2^ΔZZ^*mutant flies, where single males were paired with two or three virgin *w^1118^*females, and fertility was scored by the presence or absence of larvae in the vial after seven days. *orb2^ΔZZ^* mutants were completely sterile both as *orb2^ΔZZ^* homozygotes and when heterozygous with *orb2* deletion alleles (Figure 3a). To assess whether mature sperm were produced, we examined the seminal vesicles. Wild-type seminal vesicles from *w^1118^* males were large and full of mature sperm, as indicated by the needle-shaped nuclei of the mature sperm (Figure 3b). For all three *orb2^ΔZZ^* mutant genotypes, the seminal vesicles were small and empty, as indicated by the lack of sperm heads (Figure 3b).

**Figure 3.**
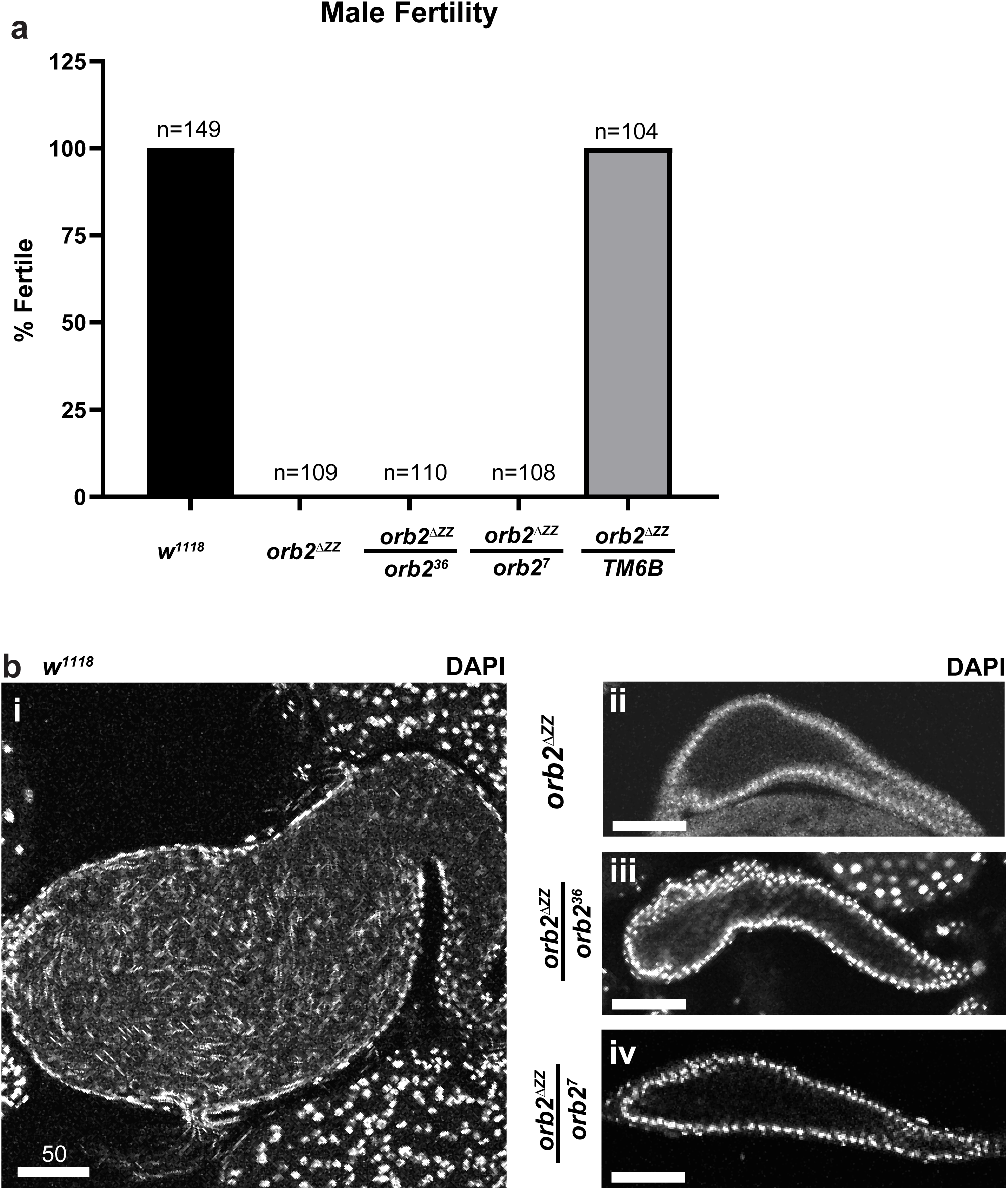
***orb2^ΔZZ^* males are sterile and have empty seminal vesicles.** (a) Bar graph showing the results of the male fertility assay, presented as a percentage. b-e) Laser-scanning confocal micrographs of whole adult testes stained with DAPI (white) zoomed in on the seminal vesicles. In *w^1118^* testes, the seminal vesicle is large and full of mature sperm, which can be identified by the needle-shaped nuclei; the seminal vesicle of *orb2^ΔZZ^* flies is empty and lacks mature sperm. Scale bars represent 50μm.

**Figure 4:**
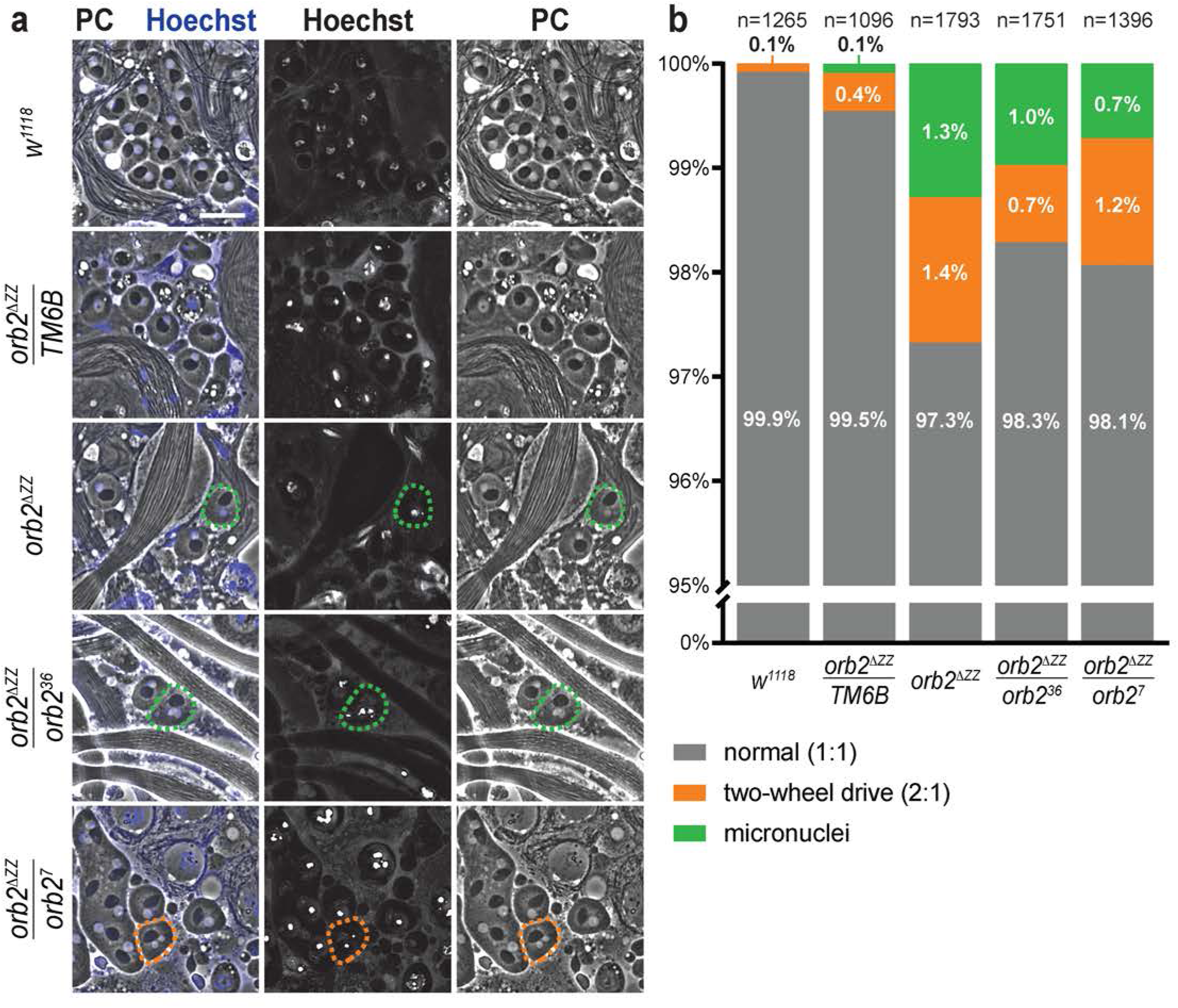
***orb2^1′ZZ^* males show defects in meiosis.** (a) Phase contrast (PC) and epifluorescence micrographs with Hoechst staining (blue) of squashed adult testes. Orange cell outline identifies a post-meiotic spermatid with a 2:1 nucleus to Nebenkern ratio (also called a “two-wheel drive” phenotype); green cell outline identifies a post-meiotic spermatid with micronuclei. Scale bar represents 20μm. b) Bar graph depicting the frequency of abnormal nucleus to Nebenkern ratios in post-meiotic spermatids. Grey represents the percentage of post-meiotic spermatids with a 1:1 nucleus to Nebenkern ratio; orange represents percentage with a 2:1 nucleus to Nebenkern ratio; green represents percentage with micronuclei.

### The ZZ domain plays a role in meiosis

Successful completion of meiosis results in 64 spermatids which have a light nucleus in phase contrast microscopy (also identifiable by positive Hoechst staining) that is paired with a dark mitochondrial Nebenkern. Spermatocytes in *orb2^36^* mutants do not complete meiosis, the Nebenkern is poorly contrasted and, occasionally, fragmented (XU *et al*. 2012).

To assess defects in meiosis upon loss of the ZZ domain, we produced and examined testis squashes (using phase contrast microscopy and staining with Hoechst) from four different genotypes: pre-injected *w^1118^* host strain that was used for CRISPR, homozygous *orb2^ΔZZ^*, and the two heterozygous genotypes *orb2^ΔZZ^*/*orb2^36^*and *orb2^ΔZZ^*/*orb2^7^* (Figure 4). 99.9% and 99.5% of post-meiotic spermatids in *w^1118^* and *orb2^ΔZZ^*/*TM6B* heterozygous controls, respectively, exhibited a nucleus to Nebenkern ratio of 1:1. In *orb2^ΔZZ^*homozygotes 2.7% of the spermatids either had a 2:1 ratio of nuclei to Nebenkern (also known as a “two-wheel drive” phenotype) or several micronuclei instead of a single nucleus. In the two hemizyous genotypes, these defects accounted for 1.7% and 1.9% of nuclei in *orb2^ΔZZ^*/*orb2^36^*and *orb2^ΔZZ^*/*orb2^7^*, respectively.

### Spermatid individualization fails in orb2^ΔZZ^ mutants

We next assessed whether *orb2^ΔZZ^* mutants exhibited defects later in spermatogenesis, notably during spermatocyte individualization, a process defective in null *orb2* mutants (XU *et al*. 2012). We stained for actin and DNA in 3-day old testes to examine the nuclear bundles and individualization complexes (IC, which contain investment cones that are marked by F-actin) formation and progression (diagrammed in Figure 1). In wild-type testes, the spermatid nuclei condense and assemble into tight bundles during elongation (Figure 5ai’). Once elongation is complete, the process of individualization begins with investment cones rich in filamentous actin that gather around each needle-shaped nucleus (Figure 5ai”). The actin cones bundle together to form the IC and travel synchronously away from the nuclei (Figure 5aii), a process that ensheathes each flagellar axoneme with its own plasma membrane and pushes the excess cytoplasm into a cystic bulge (CB) (Figure 5aiii).

**Figure 5.**
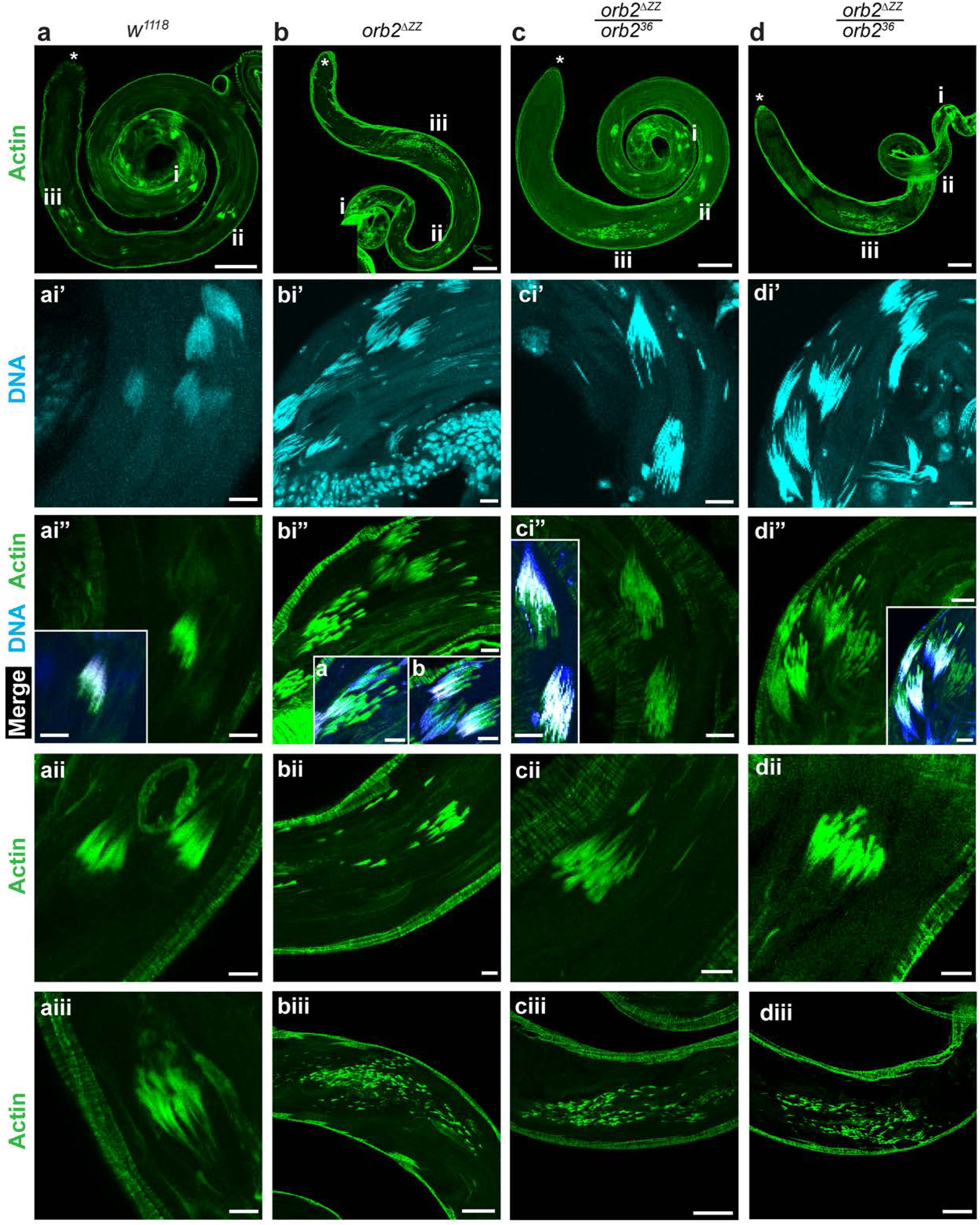
Actin cones in the individualization complex are scattered in *orb2^ΔZZ^* testes. Laser-scanning confocal micrographs of whole adult testes stained with phalloidin-FITC (green) and DAPI (blue) showing the organization of actin cones in four genotypes. a) *w^1118^*; b) *orb2^ΔZZ^*; c) *orb2^ΔZZ^*/*orb2^36^* testis; d) *orb2^ΔZZ^*/*orb2^7^* testis. Asterisk (*) indicates the apical tip of testis; scale bars represent 50μm for the top row of panels. Region i shows early individualization, scale bars represent 10μm: panel i’ shows DNA staining of nuclei organized into nuclear bundles; panel i” shows actin staining of individualization cones forming around each nucleus, inset shows merged image of i’ and i”, where blue is DNA, green is Actin, and white is the overlap; scale bars represent 10μm. Region ii depicts ICs for cysts mid-individualization; in *w^1118^*, *orb2^ΔZZ^*/*orb2^36^*, and *orb2^ΔZZ^*/*orb2^7^*, these ICs appear mostly intact, but *orb2^ΔZZ^* ICs are already dissociating and becoming scattered; scale bars represent 10μm. Region iii depicts ICs for cysts in late individualization; all three mutant genotypes have extremely scattered actin cones at this stage; scale bar in Aiii represents 10μm, and scale bars in Biii-Diii represent 50μm.

In *orb2^ΔZZ^* homozygous, *orb2^ΔZZ^*/*orb2^36^*, and *orb2^ΔZZ^*/*orb2^7^*testes, multiple nuclei were present outside of bundles; fragmenting bundles could also be observed (Figure 5b-d, panels i’). At the early stages of individualization, actin cones assembled around each nucleus as in wild type (Figure 5a, panel ii). However, in the mutants, as the ICs migrated away from the sperm head, actin cones became extremely scattered (Figure 5b-d panels ii and iii), and none of the ICs remained intact in any of the testes examined (*orb2^ΔZZ^*n = 17, *orb2^ΔZZ^*/*orb2^36^* n = 15, *orb2^ΔZZ^*/*orb2^7^*n = 13, *w^1118^* n = 19). Scattering of the actin cones began earlier in *orb2^ΔZZ^* testes compared to *orb2^ΔZZ^*/*orb2^36^*and *orb2^ΔZZ^*/*orb2^7^* (Figure 5b-d panels ii). In all three mutant genotypes, cysts in the late stages of individualization contained extremely scattered actin cones, spanning a large portion of each testis (Figure 5b-d, panels iii).

We conclude that individualization fails in *orb2^ΔZZ^* testes. This failure is likely to be the cause of male sterility that we observed in mutant adult males.

### The ZZ domain is required for enrichment of IMP, ORB and SOTI proteins at the distal tip of elongating spermatids

In elongating spermatids, proteins such as IMP, ORB, and SOTI are enriched towards the distal end (i.e., opposite end of the spermatid to where the nuclei reside) and SOTI is required for spermatid individualization (BARREAU *et al*. 2008; FABRIZIO *et al*. 2008; KAPLAN *et al*. 2010; GILMUTDINOV *et al*. 2021). We found that deletion of the ZZ domain resulted in weaker enrichment of all three of these proteins at the distal end with the proteins more dispersed along the spermatids (Figures 6a-d for IMP, 7a-d for ORB and 8a-d for SOTI). Furthermore, loss of the ZZ domain resulted in a ‘flared paintbrush’ appearance of the distal end compared to the rounded and compact shape in wild type. Since the ORB2ΔZZ protein distribution is similar to full-length ORB2 (Figure 2), these results suggest that deletion of the ZZ domain disrupts the organization and polarization of the distal portion of spermatid cysts leading to delocalization of a subset of IMP, ORB and SOTI proteins.

**Figure 6.**
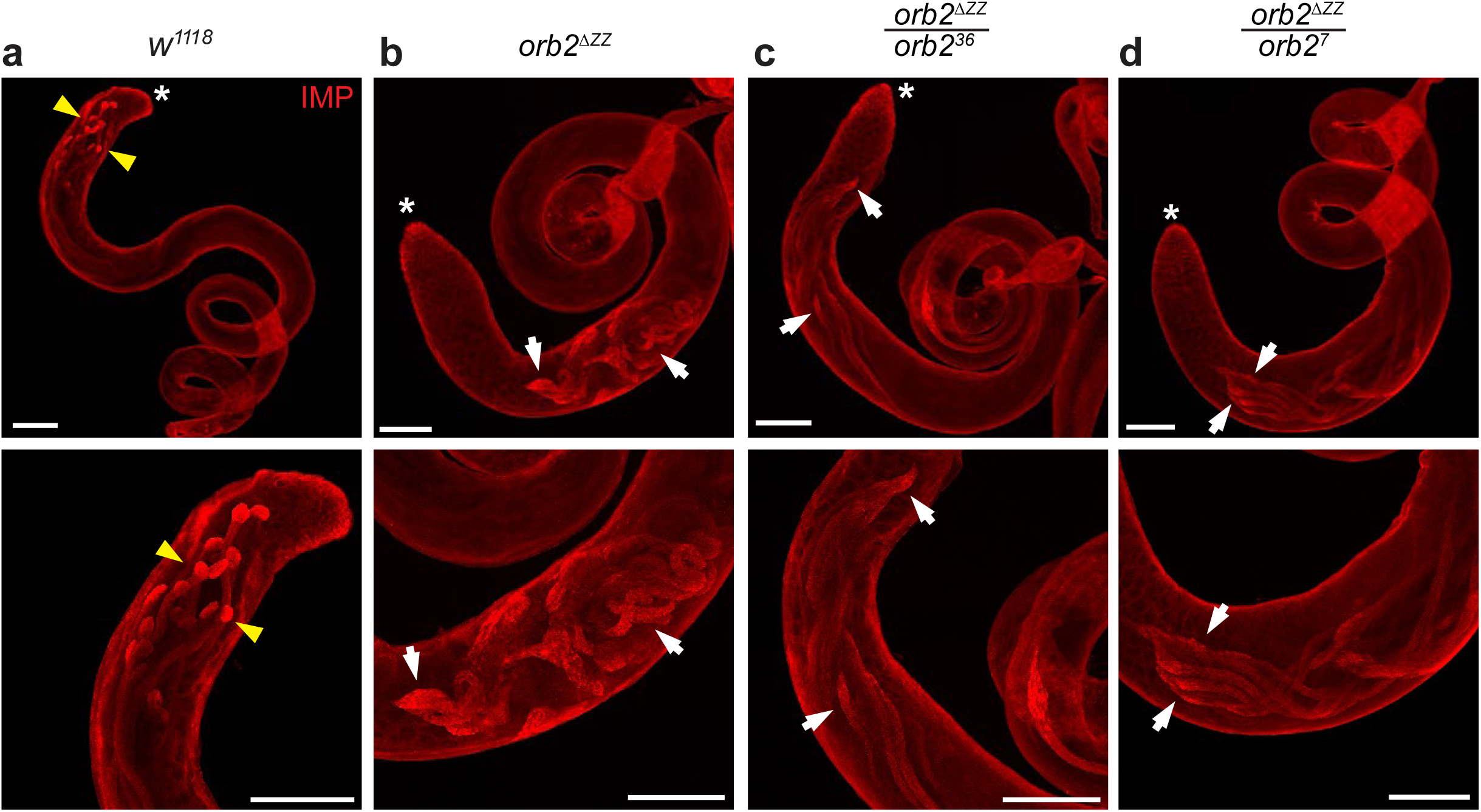
**Spatial distribution of IMP protein is disrupted in *orb2****^ΔZZ^* **testes.** Laser-scanning confocal micrographs of whole adult testes stained with anti-IMP antibody. a) *w^1118^*; b) *orb2^ΔZZ^*; c) *orb2^ΔZZ^*/*orb2^36^*; d) *orb2^ΔZZ^*/*orb2^7^*. Asterisk (*) indicates the apical tip of the testis; scale bars represent 100μm; yellow arrowheads indicate localized IMP protein, concentrated at the distal ends of cysts of elongating spermatids; white arrows indicate abnormal localization and/or expression patterns in mutant testes.

**Figure 7.**
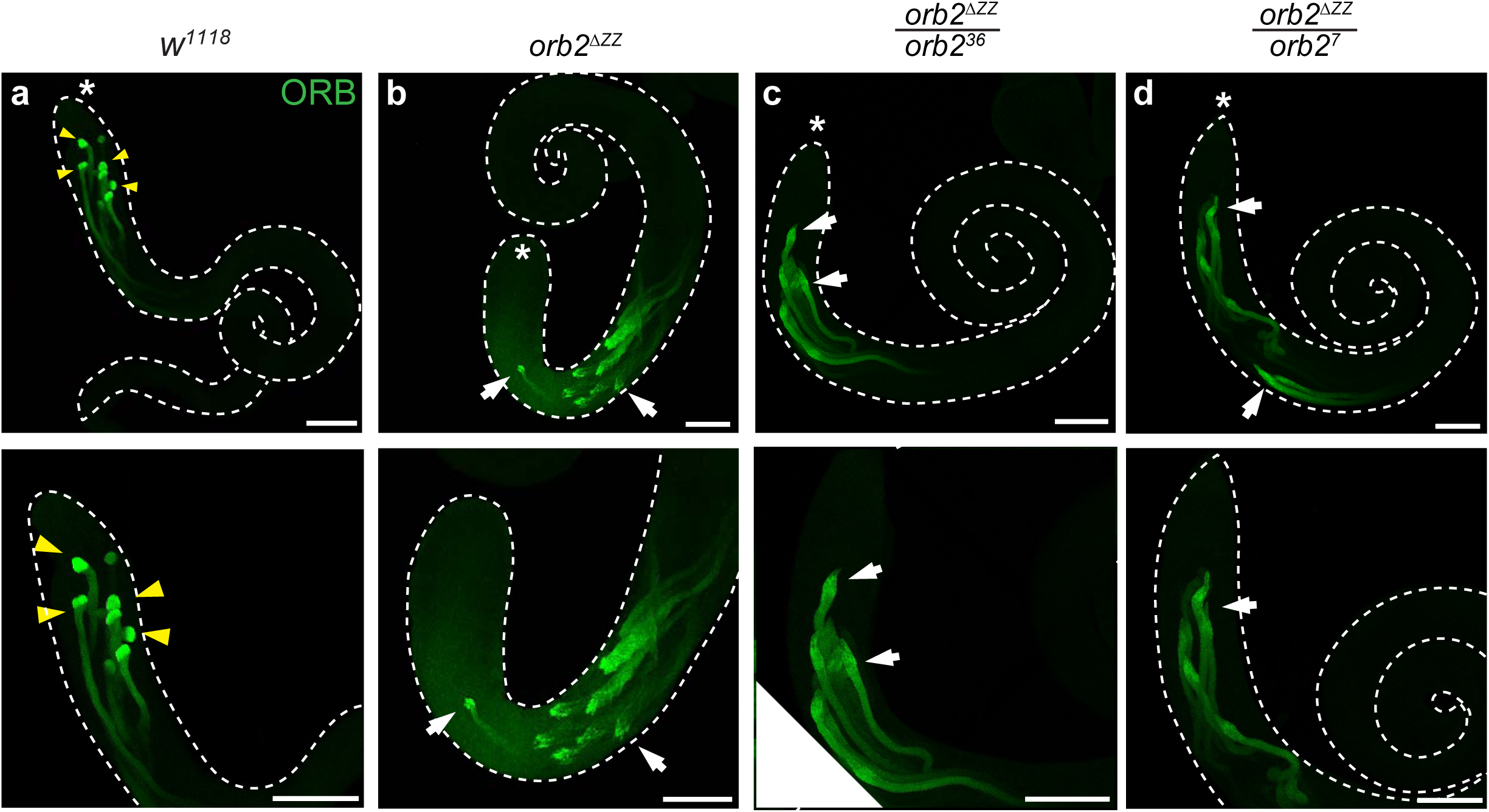
**Spatial distribution of ORB protein is disrupted in *orb2****^ΔZZ^* **testes.** Laser-scanning confocal micrographs of whole adult testes stained with anti-ORB antibody. a) *w^1118^*; b) *orb2^ΔZZ^*; c) *orb2^ΔZZ^*/*orb2^36^*; d) *orb2^ΔZZ^*/*orb2^7^*. Asterisk (*) indicates the apical tip of the testis; scale bars represent 100μm; yellow arrowheads indicate localized ORB protein, concentrated at the distal ends of cysts of elongating spermatids; white arrows indicate abnormal localization and/or expression patterns in mutant testes.

**Figure 8.**
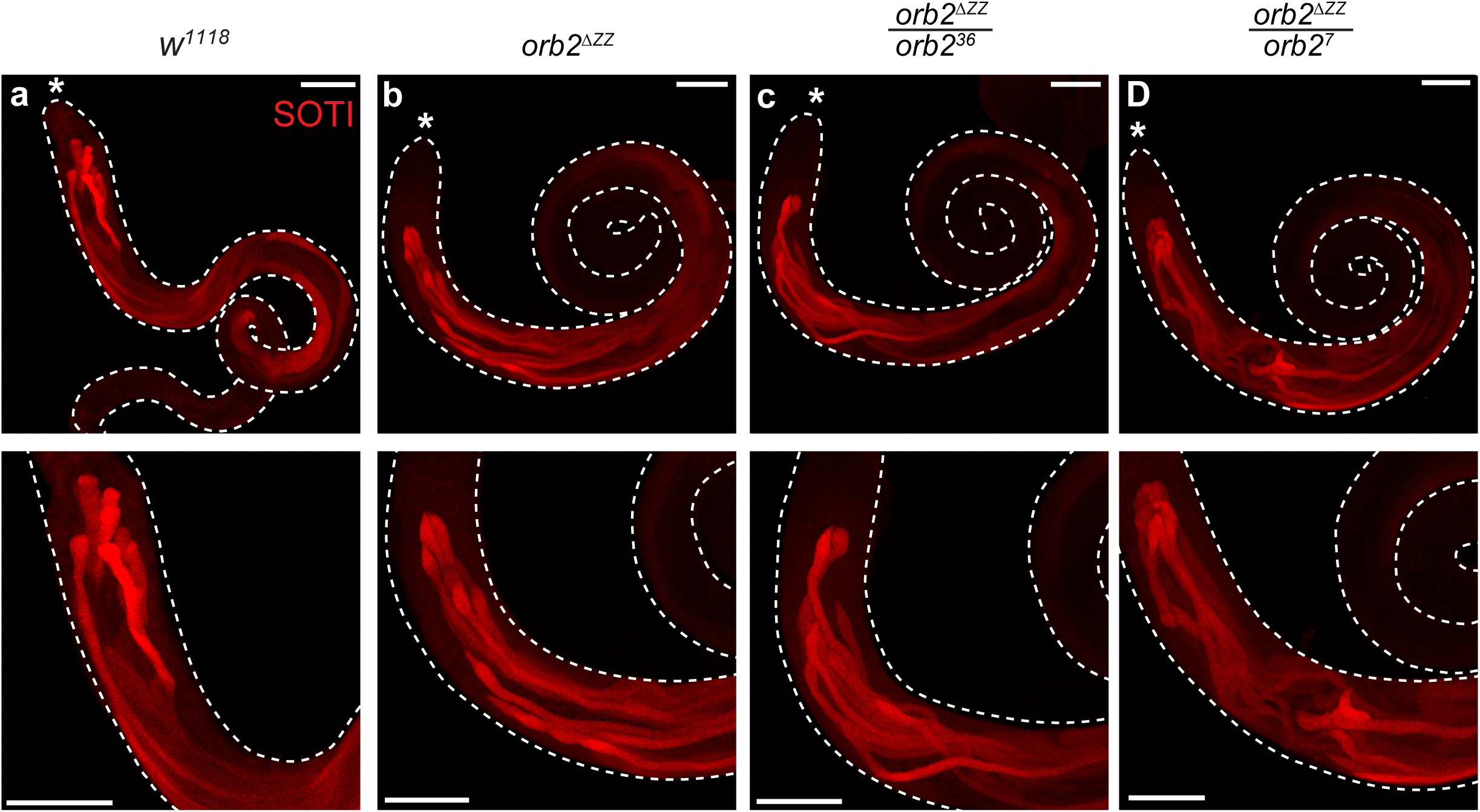
**Spatial distribution of SOTI protein is disrupted in *orb2****^ΔZZ^* **testes.** Laser-scanning confocal micrographs of whole adult testes stained with anti-SOTI antibody. a) *w^1118^*; b) *orb2^ΔZZ^*; c) *orb2^ΔZZ^*/*orb2^36^*; d) *orb2^ΔZZ^*/*orb2^7^*. Asterisk (*) indicates the apical tip of the testis; scale bars represent 100μm.

**Figure 9:**
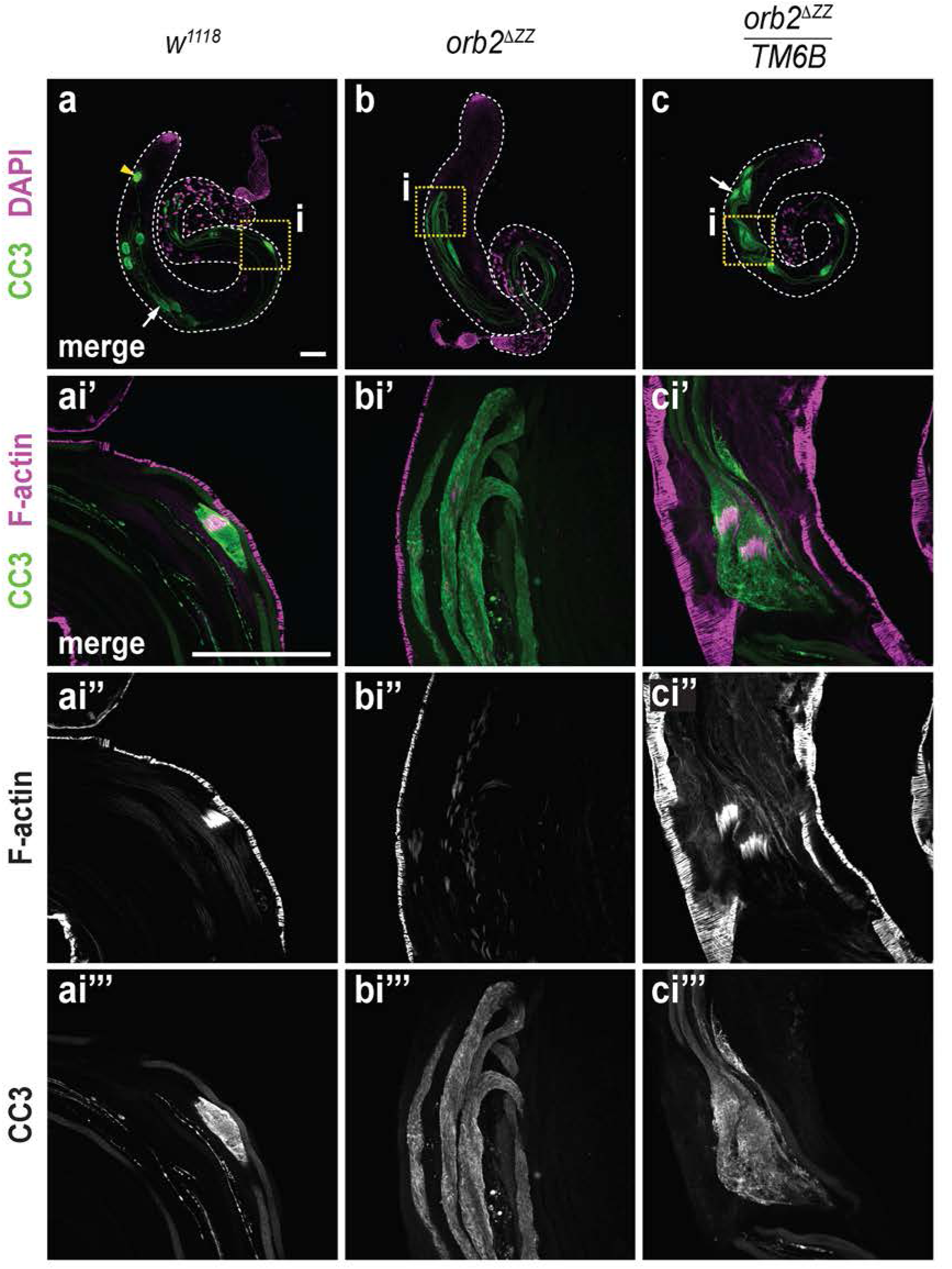
The CC3 protein gradient is disrupted and CC3 is overexpressed in *orb2^1′ZZ^* testes. Laser-scanning confocal micrographs of whole adult testes stained with anti-CC3 (green) and rhodamine phalloidin (magenta). a) *w^1118^*; b) *orb2^1′ZZ^*; c) *orb2^1′ZZ^*/*TM6B.* White arrows indicate examples of cystic bulges; yellow arrowhead indicates example of a waste bag. Scale bar represents 100μm.

### The CC3 gradient is disrupted in orb2^ΔZZ^ mutants

The SOTI protein gradient has been shown to be important for the successful completion of individualization. Specifically, the SOTI gradient regulates an opposing gradient of activated (also called cleaved) caspase-3 (KAPLAN *et al*. 2010). Cleaved caspase-3 (CC3) is localized in a pattern opposite to that of SOTI protein because SOTI represses CC3 activity via a Cullin-3-based E3 ubiquitin ligase complex. In wild-type cysts, low levels of CC3 are at the distal end of the spermatid cyst (Figure 9a) and activity increases towards the spermatid nuclei, with CC3 concentrated at the IC. This concentration of CC3 activity is pushed down the length of the cyst along with cystic bulge by the IC as it travels away from the nuclei, eventually being discarded inside the waste bag.

Since the SOTI gradient is disrupted in the testes of *orb2^ΔZZ^* mutants, we asked whether the CC3 gradient was also affected. We found that the CC3 gradient was altered such that CC3 protein was distributed along the length of the cyst (Figure 9b, c). In addition, we found that the cystic bulges in *orb2^ΔZZ^* testes were wider and longer compared to wild-type testes and waste bags are completely absent in *orb2^ΔZZ^*testes.

Together, our data are consistent with a model in which the ZZ domain is required for organization of the distal region of the spermatid cyst and establishment of the SOTI gradient, which in turn regulates CC3 production and successful spermatid individualization.

## Discussion

Here, we have shown that the ZZ domain of ORB2 is required for male fertility and is involved in at least two aspects of spermatogenesis: meiosis of spermatocytes to produce spermatids and, subsequently, spermatid individualization to produce mature sperm. We observed defects in meiosis in *orb2^ΔZZ^* mutants, both when homozygous and when present in one copy when heterozygous with an *orb2* deletion. While these meiotic defects were of low penetrance, it is important to note that many, small nuclei could lead to the formation of numerous small actin cones (multiple cones per spermatid, i.e., many more than 64 cones per cyst) with subsequent failure of individualization and consequent male sterility. Indeed, the phenotype of *orb2^ΔZZ^*mutants resembles that of *orb2^ΔQ^* and *orb2^R^* mutants – respectively, deleting the polyQ domain or mutating the RBD – which also have low penetrance meiotic defects but a strong effect on individualization: Fragmentation and scattering of actin cones, mispatterning of key protein gradients (ORB, SOTI, and CC3), and a complete failure of individualization (XU *et al*. 2012; XU *et al*. 2014; GILMUTDINOV *et al*. 2021).

The observation that the polyQ domain is required for sperm individualization suggests that the ability of ORB2A and ORB2B to oligomerize is required for the completion of spermatogenesis (XU *et al*. 2012). The polyQ domain has been associated with translational activation of ORB2 bound transcripts (WHITE-GRINDLEY *et al*. 2014; KHAN *et al*. 2015). Our recent analysis of the role of the ZZ domain in the early embryo has shown that it is required for translational repression of ORB2’s target mRNAs (LOW *et al*. 2025); however, since ORB2B is the only isoform expressed in early embryos, that study does not shed light on the possible role of the ZZ domain in translational activation in the context of ORB2A-ORB2B hetero-oligomers. The fact that *orb2^ΔZZ^* mutants phenocopy the *orb2^ΔQ^* mutant is consistent with a role for the ZZ domain in ORB2-mediated translational activation.

We have recently shown that the ZZ domain interacts with all of the components of the 43S translation preinitiation complex (PIC) in early embryos (LOW *et al*. 2025). During cap-dependent translation initiation, the PIC is recruited to the 5‘-end of an mRNA by eIF4F, which includes eIF4A, the cap-binding protein eIF4E, and eIF4G. If ORB2’s interaction with the PIC is conserved in testes, then we speculate that translational activation by the ORB2A-ORB2B heteromers may be mediated at least in part by recruitment of the 43S PIC to target mRNAs. We recently published link between ORB2 and the testis-specific eIF4E paralog, eIF4E5, which is consistent with such a role (SHAO *et al*. 2023): ORB2 and eIF4E5 genetically interact to control spermatid cyst polarization; additionally, eIF4E5 mutants display defects in SOTI protein accumulation and defective individualization, resulting in male sterility. Whether PIC and eIF4E5 interaction is the basis for ORB2-mediated activation in the testis will be an interesting area for future study.

## Data Availability

The authors affirm that all of the data necessary for confirming the conclusions of the article are present within the article, figures and supplementary figure.

## Acknowledgements

We thank Eli Arama and Paul Macdonald for providing anti-SOTI and anti-IMP antibodies, respectively. We also thank Kimberly Lau and Paul Paroutis of the SickKids Imaging facility for assistance with imaging.

---

**Figure S1: Map of the *orb2* locus showing location and sequence of primers used in Figure 2a and b**.

